# Structural regeneration and functional recovery of the olfactory system of zebrafish following brain injury

**DOI:** 10.1101/2024.12.03.626704

**Authors:** Erika Calvo-Ochoa, Nathaniel W. Vorhees, Theodore P. Lockett, Skylar L. DeWitt, Evan A. Thomas, Abigail B. Gray, Nobuhiko Miyasaka, Yoshihiro Yoshihara, Christine A. Byrd-Jacobs

## Abstract

Olfactory dysfunction is a common outcome of brain injuries, negatively affecting quality of life. The mammalian nervous system has limited capacity for spontaneous olfactory recovery, making it challenging to study olfactory regeneration and recovery in adults. In contrast, zebrafish are an ideal model for such studies due to its extensive and lifelong regenerative abilities. In this work, we describe a model of excitotoxic injury in the olfactory bulb using quinolinic acid (QA) lesions in adult zebrafish. We observed extensive neurodegeneration in both the olfactory bulb and olfactory epithelium, including a reduction of bulbar volume, neuronal death, and impaired olfactory function. Recovery mechanisms involved tissue remodeling, cell proliferation, neurogenesis, leading to full restoration of olfactory function by 21 days. This study provides a model to further investigate the effects of excitotoxicity on olfactory dysfunction, and highlights zebrafish’s remarkable regenerative abilities, providing insights into potential therapeutic strategies for restoring olfactory function following brain injuries.

## INTRODUCTION

The sense of smell plays a crucial role in maintaining good health and well-being by enabling the perception of important nutritional, environmental, and social chemical cues necessary for survival (Firestein et al., 2001). The olfactory system, which enables the sense of smell, has remained well-conserved throughout vertebrate evolution ((Hildebrand & Shepherd, 1997; Saraiva et al., 2015).The olfactory sensory neurons (OSNs) in the olfactory epithelium (OE) detects odorants through the activation of olfactory receptors and transmit signals to the olfactory bulb (OB) (Byrd & Brunjes, 1995; Sato et al., 2005). Olfactory signals are processed through intrinsic neural circuitry in the OB, where output neurons relay these signals to higher olfactory centers for further processing (Friedrich et al., 2010; Miyasaka et al., 2009). Olfactory dysfunction is a disability that significantly reduces individuals’ quality of life (Miwa et al., 2001) and a common and early symptom of neurodegenerative diseases such as Alzheimer’s disease and Parkinson’s disease (Fullard et al., 2017; Wilson et al., 2009). Brain injury or trauma commonly results in olfactory impairment, with up to 68% of traumatic brain injuries leading to some degree of olfactory loss (Schofield et al., 2014).

The OE is known for its constitutive cell turnover, driven by stem cells that replace lost sensory neurons (Schwob et al., 2017) The OB also exhibits life-long neurogenesis (Lledo & Valley, 2016) with stem cells in the subventricular zone generating neural precursor cells that migrate to the OB and become granule and periglomerular cells (Lois & Alvarez-Buylla, 1994; Takahashi et al., 2018). Even with extensive plasticity in the OE and OB, in mammals, most individuals with post-injury olfactory dysfunction do not experience spontaneous functional recovery in subsequent years (Doty et al., 1997; Mattar & El Adle, 2020; Reden et al., 2006). This discrepancy is thought to arise from improper neural integration of OSNs into the OB (Butler et al., 1984) and from incomplete OB reorganization following damage (Graziadei & Samanen, 1980; Monti-Graziadei & Graziadei, 1992). Thus, studying mammalian models can provide only a partial understanding of olfactory regeneration and recovery processes (Richardson et al., 2007). To address this issue, we used zebrafish (*Danio rerio*), an organism with widespread and lifelong neuronal regenerative abilities (Calvo-Ochoa et al., 2020; Hentig & Byrd-Jacobs, 2016; Hinsch & Zupanc, 2007; Zupanc, 2008). An important process underlying this regenerative capacity is the generation of new neurons (i.e., neurogenesis), which occurs constitutively across 16 unique neurogenic niches in the zebrafish brain (Adolf et al., 2006; Marz et al., 2010).

The olfactory system of zebrafish exhibits outstanding neuroplasticity, with significant regenerative and repair capabilities following direct injury to the OE. This includes complete repair and regeneration of the OE itself (Hentig & Byrd-Jacobs, 2016; Iqbal & Byrd-Jacobs, 2010; Kocagöz et al., 2022) and the OB (Paskin et al., 2011; White et al., 2015), highlighting its remarkable ability to recover following peripheral injuries.

While the neurodegeneration and recovery processes following OE damage are well-documented, the impact of OB injuries on the overall structure and function of the olfactory system, including whether it prompts repair and regeneration, is less explored. Motivated by these questions, we developed a novel OB injury model using the excitotoxic drug quinolinic acid (QA) to investigate the effects of OB damage on the morphology, function, and recovery of the olfactory system. We sought to investigate bulbar excitotoxicity in zebrafish, as it may serve as a mechanistic link between brain injury and olfactory loss (Marin et al., 2022; Marin et al., 2017).

Here, we provide a comprehensive and integrated characterization of the structural and functional consequences of an excitotoxic lesion to the OB and the subsequent processes that lead to repair and recovery by 21 days postlesion (dpl). Our results show that OB lesions lead to extensive neurodegeneration throughout the olfactory system, including severe structural changes in the OB, altered olfactory glomerular morphology, and loss of OSNs in the OE, resulting in functional impairment. Notably, by 21 days, the lesion-induced altered morphology and functional deficits were largely restored. Additionally, OB lesions induced proliferation and neurogenesis in the OE, the OB, and the telencephalon, processes that contribute to recovery. We also report increased neuroinflammation, which has been associated with proliferation and neural recovery in other brain regions (Kyritsis et al., 2012).

## RESULTS

### QA injections induce neurodegeneration and recovery in the olfactory bulb

Our objective was to investigate the structural recovery and remodeling responses of the zebrafish OB following injury. We developed a novel bulbar lesion using quinolinic acid (QA), an NMDA receptor agonist commonly used in models of excitotoxic damage and neurodegenerative disorders (Lugo-Huitrón et al., 2013; Skaggs et al., 2014). In our lesion paradigm, we injected the right OB (referred to as ipsilateral or ipsi, indicated in yellow in Fig. 1B) while leaving the left side as an internal control (contralateral or contra, indicated in blue in Fig. 1B). To assess bulbar structural damage and neural loss, we examined coronal sections immunostained with HuC/D, a pan-neuronal marker. At 1 dpl, the ipsilateral bulb showed a noticeable reduction in size, while the contralateral side was comparable to the unlesioned control (Fig. 1A left and middle panels. In the 21 dpl groups, the ipsilateral bulb retained its original appearance (Fig. 1A right panel). In addition, there was a noticeable disruption of the two innermost OB laminae (Fig. 1A, A’). The glomerular layer (GL) comprises olfactory axon termini, dendrites and somata of output neurons (i.e., mitral and ruffed cells; (Fuller et al., 2006; Sato et al., 2005), and interneurons (i.e., periglomerular cells (S. T. Bundschuh et al., 2012; Byrd & Brunjes, 1995). The internal cell layer (ICL) consists somata of granule cell interneurons ((Byrd & Brunjes, 1995; Edwards & Michel, 2003; Zhu et al., 2013)). High magnification images revealed changes in neuronal distribution and OB morphology, as well as alterations in cell nuclei (DAPI+) distribution in both layers in the ipsilateral side of the 1 dpl group. At 21 dpl, some cell reorganization of cell nuclei was observed (Fig. 1A’ upper and lower panels).

**Figure 1.**
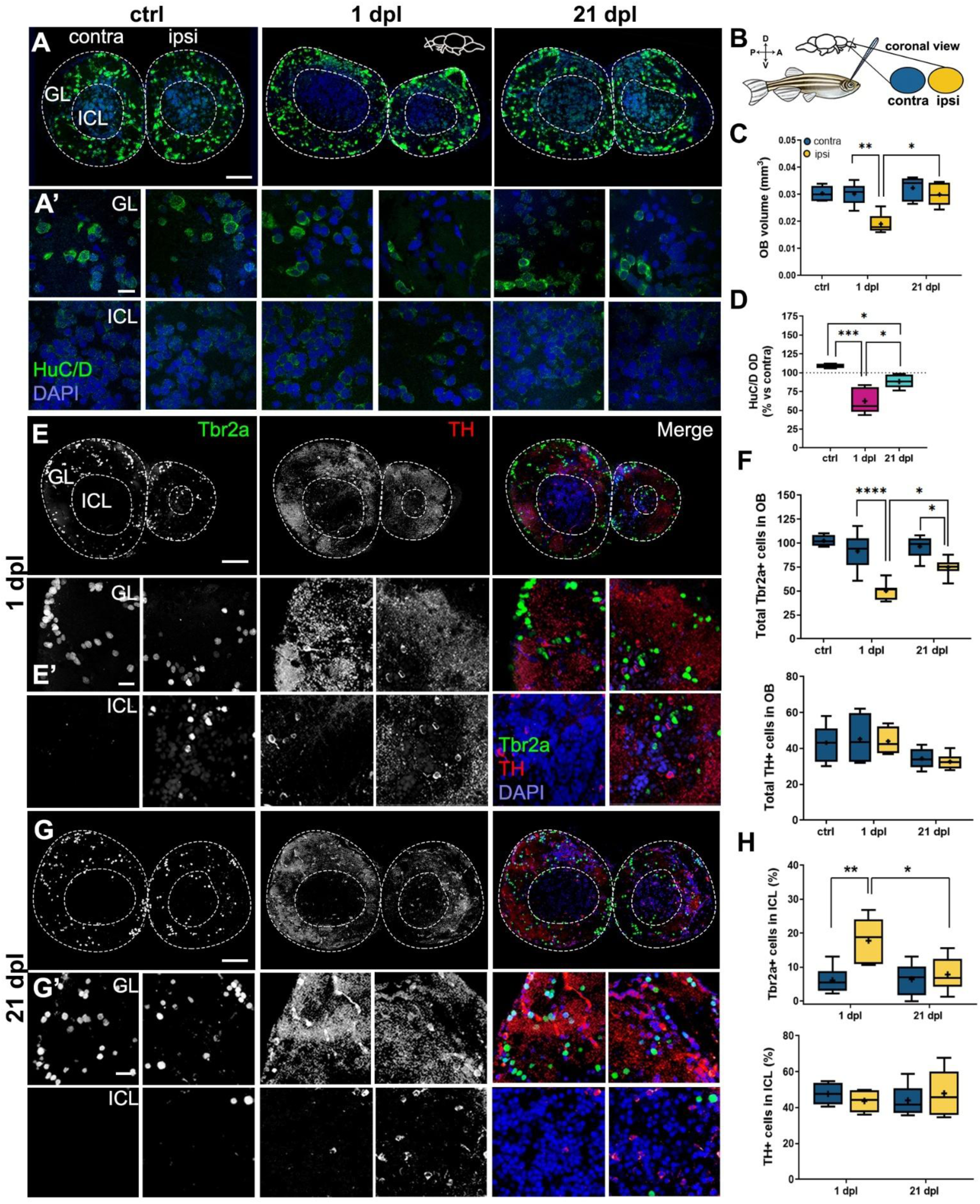
Effects of an excitotoxic lesion on the olfactory bulb at 1-day post-lesion (dpl) and 21 dpl. A) and A’) HuC/D immunohistochemistry of coronal sections of the OB from unlesioned controls (left panels), 1-dpl (middle panels) and 21-dpl fish (right panels). Lower panels are magnified views of the glomerular layer (GL) and intracellular layer (ICL), respectively. Green: HuC/D; blue, DAPI. Scale bars: 50 µm in (A); 20 µm in (A’). B) Schematic of bulbar lesioning method with QA (quinolinic acid) injections. The contralateral side is shown in blue (left), while the ipsilateral side (right) is shown in yellow. C) Quantification of average OB volume on semi-serial H&E-stained coronal sections (n = 3-6). D) Quantification of percent change (vs. contra) in optical density (OD) of HuC/D in the OB from (A) (n = 4-7). E), E’), G) and G’) Double immunohistochemistry of Tbr2a and tyrosine hydroxylase (TH) of coronal sections of the OB from 1-dpl (E, E’) and 21-dpl fish (G, G’). Upper and lower panels in (E’) and (G’) are magnified views of the glomerular layer (GL) and intracellular layer (ICL), respectively. Green: Tbr2a; red: TH; blue, DAPI. Scale bars: 50 µm in (E) and (G); 20 µm in (E’) and (G’). F) and H) Quantification of the number of Tbr2a+ and TH+ cells in whole bulb (F) and the ICL (H) in coronal sections of the OB from (E) and (G) (n = 4-8). Box plots indicate mean (+), quartiles (boxes) and range (whiskers). One-way ANOVA, *p < 0.005, **p < 0.001, ***p = 0.0009, ****p < 0.0001.

To quantitatively assess the lesion’s extent, we calculated the bulbar volume in ipsi- and contralateral sides of coronal OB sections using the stereological Cavalieri method followed by a Gundersen coefficient of error (CE) analysis (Fig. S1). We observed significant volume reduction in the ipsilateral OB due to the QA lesion, with no effect on the contralateral OB’s volume (Fig. 1C). Remarkably, bulbar volume was restored to control levels by 21 dpl (Fig. 1C), indicating extensive histological reorganization. To assess if these bulbar changes were associated with neuronal loss, we compared the staining intensity of HuC/D. We found decreased neuronal staining on the lesioned side compared to the contralateral side (Fig. 1D). Interestingly, while neuronal density recovered by 21 dpl, HuC/D staining remained significantly different from the unlesioned controls (Fig. 1D). This partial recovery of cell density aligns with our observations in high magnification images (Fig. 1A’). Next, we sought to determine if QA specifically targets mitral cells, the major glutamatergic output neurons in the bulb (Edwards & Michel, 2003; Friedrich & Laurent, 2001). We examined immunostaining for Tbr2a, a marker for a subset of mitral cells (Miyasaka et al., 2009), and tyrosine hydroxylase (TH), a marker of dopaminergic periglomerular interneurons (Sebastian T Bundschuh et al., 2012; Yamamoto et al., 2010), respectively. We confirmed that QA lesions decreased the number of Tbr2a+ mitral cells while sparing TH+ periglomerular neurons (Fig. 1E, F).

We also examined the localization of these neuronal subtypes within the bulbar laminae, noting disturbances in neuronal localization (Fig. 1E, E’). Anti-TH staining delineated the location of the GL, where apical dendrites of periglomerular cells are found. Mitral cell somata (Tbr2a+) are also located within this layer (Fig. 1E’ upper panel). At 1 dpl, we observed that the somata of large neurons (i.e., mitral and periglomerular cells) were redistributed to the ICL on the lesioned side (Fig. 1E’ lower panel). Additionally, some TH+ puncta (i.e., dendrites) were redistributed to the ICL (Fig. 1E’ lower panel), indicating significant disruption of cellular localization within bulbar laminae.

By 21 dpl, we observed substantial reorganization and remodeling of bulbar laminae, with the location of mitral cell somata and periglomerular cell somata and dendrites resembling that of the contralateral side (Fig. 1 G, G’). Interestingly, we found a partial recovery of mitral cells (Fig. 1F). Tbr2a+ mitral cells in the ipsilateral side of 21-dpl OB were significantly different from both the 1-dpl group and the unlesioned controls (Fig. 1F). While the total number of TH+ cells was not affected, TH+ cell counts in the ICL were significantly different from both the 1 dpl group and the unlesioned controls (Fig. 1H). Collectively, our results show that unilateral QA injections caused extensive neurodegeneration and histological disruption in the OB, specifically targeting Tbr2a+ mitral cells, while the contralateral bulb remained intact. Notably, structural recovery of the lesioned side occurred by 21 dpl, with laminar reorganization and partial restoration of neuron density.

### QA lesions cause disruption of olfactory glomeruli and olfactory epithelium followed by recovery

Olfactory glomeruli are spheroidal structures where afferent axonal fibers from the OE and dendrites of bulbar neurons form synaptic connections. The shape and location of glomeruli in the OB are stereotypical and well-characterized (Baier & Korsching, 1994; Braubach et al., 2012). Given that OB lesions disrupt laminar organization and cause neuronal loss, we aimed to determine if QA lesions affect the morphology and integrity of olfactory glomeruli. We used whole-mount brain preparations to conduct qualitative immunohistochemical analysis with three markers: anti-calretinin, for a subset of ciliated and microvillous OSNs innervating the dorsal, dorsolateral, ventromedial, and lateral clusters; anti-Gαolf for ciliated OSNs innervating the dorsal, dorsolateral, and ventromedial clusters; and anti-SV2 (synaptic vesicle protein 2), which highlights glomerular structure as it labels pre-synaptic terminals (Braubach et al., 2012). The localization of these markers in the OB is illustrated in Fig. S2A. We examined both the dorsal and ventral sides of the OB to characterize the major glomerular clusters in these regions.

On the dorsal side of the lesioned bulb, we observed disruption in axonal and glomerular morphology at 1 dpl (Fig. 2A). Higher magnification images revealed substantial axonal disorganization in anti-calretinin labeled fibers and even more severed disruption of anti-Gαolf stained axons, though the general location of both clusters remained unchanged (Fig. 2A). We noted disturbances in the organization and shape of axonal bundles within the glomeruli compared to the contralateral side (Fig. 2A’, A’’, arrowheads). Additionally, defasciculation of axonal termini was observed in some Gαolf+ smaller glomeruli in the dorsal cluster (Fig. 2A’’, arrowheads). These disruptions were accompanied by changes in anti-SV2 staining, which appeared to be disorganized in some dorsolateral and dorsal clusters (Fig. 2A’, A’’, arrowheads). Interestingly, calretinin+ ventromedial glomeruli on the ventral side of the lesioned bulbs showed no major structural or morphological changes (Figs. S2B, C).

**Figure 2.**
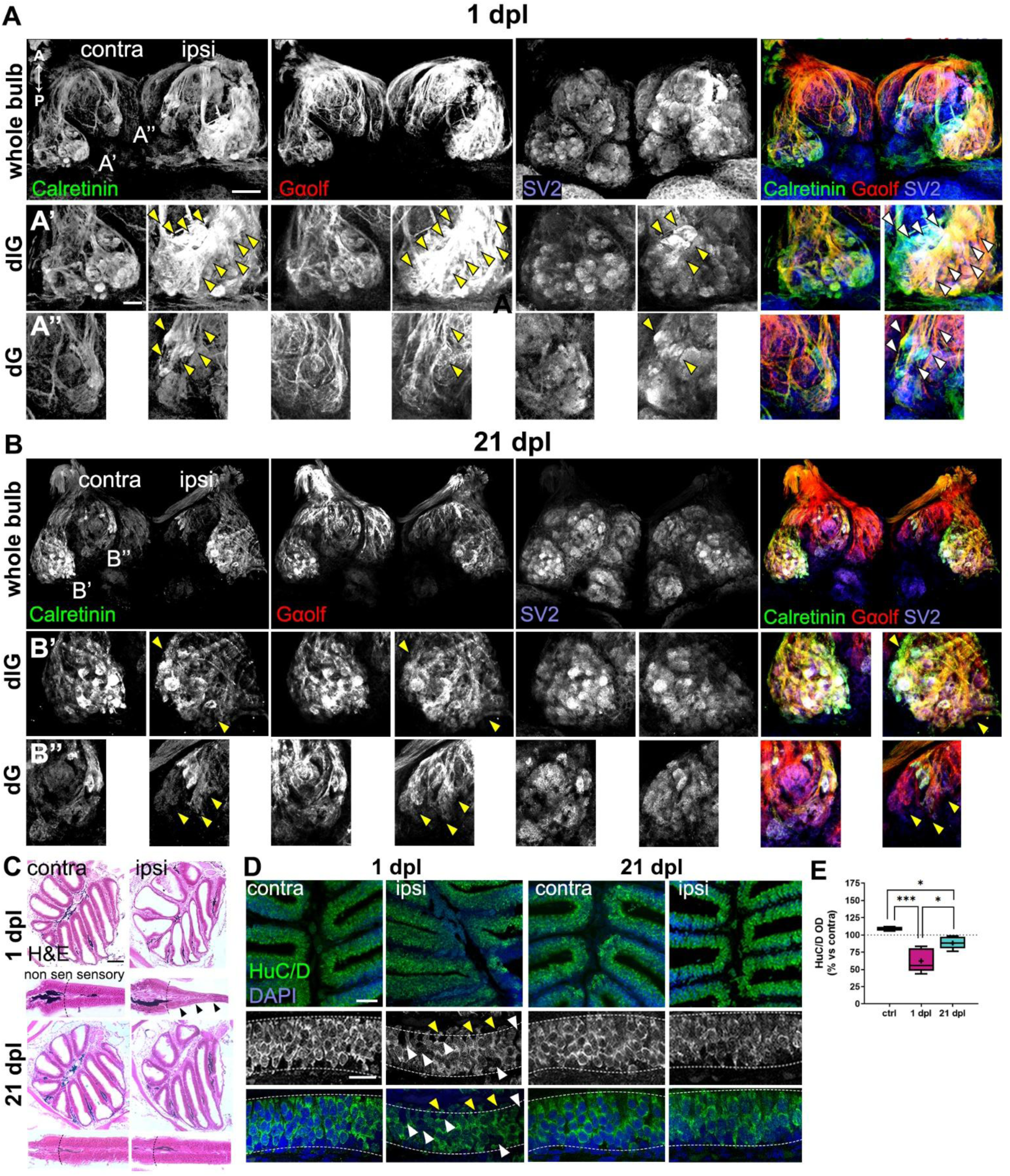
Effects of QA lesion on olfactory glomerular morphology and olfactory epithelium structure at 1 dpl and 21 dpl. A) and B) Dorsal views of whole-mount OB of 1-dpl (A) and 21-dpl (B) groups immunostained against calretinin (left panels), Gαolf (middle panels), and SV2 (synaptic vesicle protein 2, right panels). Scale bar: 100 µm. Abbreviations for glomerular clusters are: dG, dorsal, and dlG, dorsolateral. A’) and A”) Magnified views of the dorsolateral and dorsal clusters from A). B’) and B”) Magnified views of the dorsolateral and dorsal clusters from (B). Axonal defasciculation and disorganization are indicated with arrowheads. Green: calretinin; red: Gαolf; blue, SV2. Scale bar: 20 µm. C) Hematoxylin and Eosin (H&E) staining of the whole olfactory organ of 1-dpl (upper panels) and 21-dpl (lower panels) groups. Panels below each picture show a magnified view of the olfactory epithelium, in which the boundary between sensory and non-sensory regions is indicated (dashed lines). Degeneration of the sensory region is indicated with black arrowheads. Scale bars: 100 µm upper panels; 20 µm lower panels. D) HuC/D immunohistochemistry of OE sections of 1-dpl (left panels) and 21-dpl (right panels) groups. Lower panels are magnified views of the sensory olfactory epithelium. OSN degeneration and disorganization of the epithelial structure is indicated with white arrowheads. Disruption of apical dendrites is indicated with yellow arrowheads. Green: HuC/D; blue, DAPI. Scale bars: 50 µm upper panels; 20 µm lower panels. E) Quantification of percent change (vs. contra) in optical density (OD) of HuC/D in the OE from (D) (n = 10). Box plots indicate mean (+), quartiles (boxes) and range (whiskers). One-way ANOVA, *p < 0.005, **p < 0.001. A schematic of the dorsal and ventral glomerular clusters assessed can be found in supplemental figure 4.

By 21 dpl, glomerular clusters had mostly reorganized and recovered their normal organization and morphology (Fig. 2B). Although some axons in the dorsolateral cluster remained disorganized, the structure of individual glomeruli appeared closer to the unlesioned side than to the lesioned bulbs at 1 dpl (Fig. 2B, B”). On the other hand, the dorsal cluster glomeruli still appeared defasciculated and incomplete (Fig. 2B’, B’’).

To investigate if extensive axonal defasciculation and neuronal loss in the OB led to peripheral OSN loss due to retrograde degeneration (Graziadei & Monti Graziadei, 1980; Holcomb et al., 1995) we examined the histological appearance of the entire olfactory organ in Hematoxylin-Eosin (H&E) stained sections (Fig. 2G). We observed noticeable thinning of the sensory region of the OE on the ipsilateral side compared to the control side (Fig. 2C, lower panels). There was no sign of widespread degeneration of the entire olfactory organ, similar to what has been described in direct OE damage models (Iqbal & Byrd-Jacobs, 2010; Kocagöz et al., 2022). Anti-HuC/D labeling revealed disorganization of the sensory portion of lamellae, (Fig. 2D, left panels, white arrowheads) and disturbances in OSN morphology, including changes in cell body shape and disruption of apical dendrites (Fig. 2D, left panels, yellow arrowheads). We also observed a significant reduction in HuC/D staining on the ipsilateral side compared to controls, indicative of OSN loss (Fig. 2E). Remarkably, by 21 dpl, the olfactory organ had recovered its pre-lesioned structure and appearance as well as HuC/D staining levels (Fig. 2E), with restored OSN organization and morphology (Fig. 2G, right panels), indicating OSN turnover and epithelial reorganization.

### QA lesions cause impaired olfactory function, which is fully restored by 21 dpl

Given the extent of post-lesion degeneration and subsequent structural recovery throughout the olfactory system, we predicted that olfaction would be impaired initially but would recover by 21 dpl. To test this, we examined olfactory-mediated behavioral responses to three odorants with physiological relevance: alanine (a food cue), a combination of urea and taurocholic acid (TCA) (kinship cues), and cadaverine (an aversive cue). Alanine activates microvillous OSNs, whereas TCA, urea, and cadaverine activate ciliated OSNs (Dieris et al., 2017; Hussain et al., 2013; Koide et al., 2009; Lipschitz & Michel, 2002). The behavioral responses to these odorants are well-documented and stereotypical (Kalueff et al., 2013).

Single, acclimated fish were exposed to individual odorants for testing. We recorded swimming behaviors for 30 seconds before (pre-odorant) and 30 seconds after (post-odorant) odorant exposure and then analyzed various behavioral parameters. Prior to the trials, we confirmed that most of the odorant solution remained in half of the chamber 30 seconds after application (Fig. 3A). Control fish displayed expected behavioral responses to all tested odorants. Following alanine exposure, fish displayed a significant preference for the side where the amino acid was introduced, indicative of a robust attraction to amino acids (Fig. 3B, C; (Koide et al., 2009; Wakisaka et al., 2017). Control fish responded to a mixture of urea and TCA, chemicals present in fish excretions that elicit robust behavioral responses (Kermen et al., 2020; Paskin & Byrd-Jacobs, 2012)by swimming towards the odorant side and increasing directional changes (Fig. 3D, E). After cadaverine exposure, fish displayed stereotypical aversive behaviors, alternating between erratic swimming and freezing (Fig. 3F, (Hussain et al., 2013). Importantly, swimming speed was not altered in lesioned fish at 1 dpl or 21 dpl (Fig. S3), and we did not observe behavioral alterations during the trials, indicating that the QA lesion does not impair swimming.

**Figure 3.**
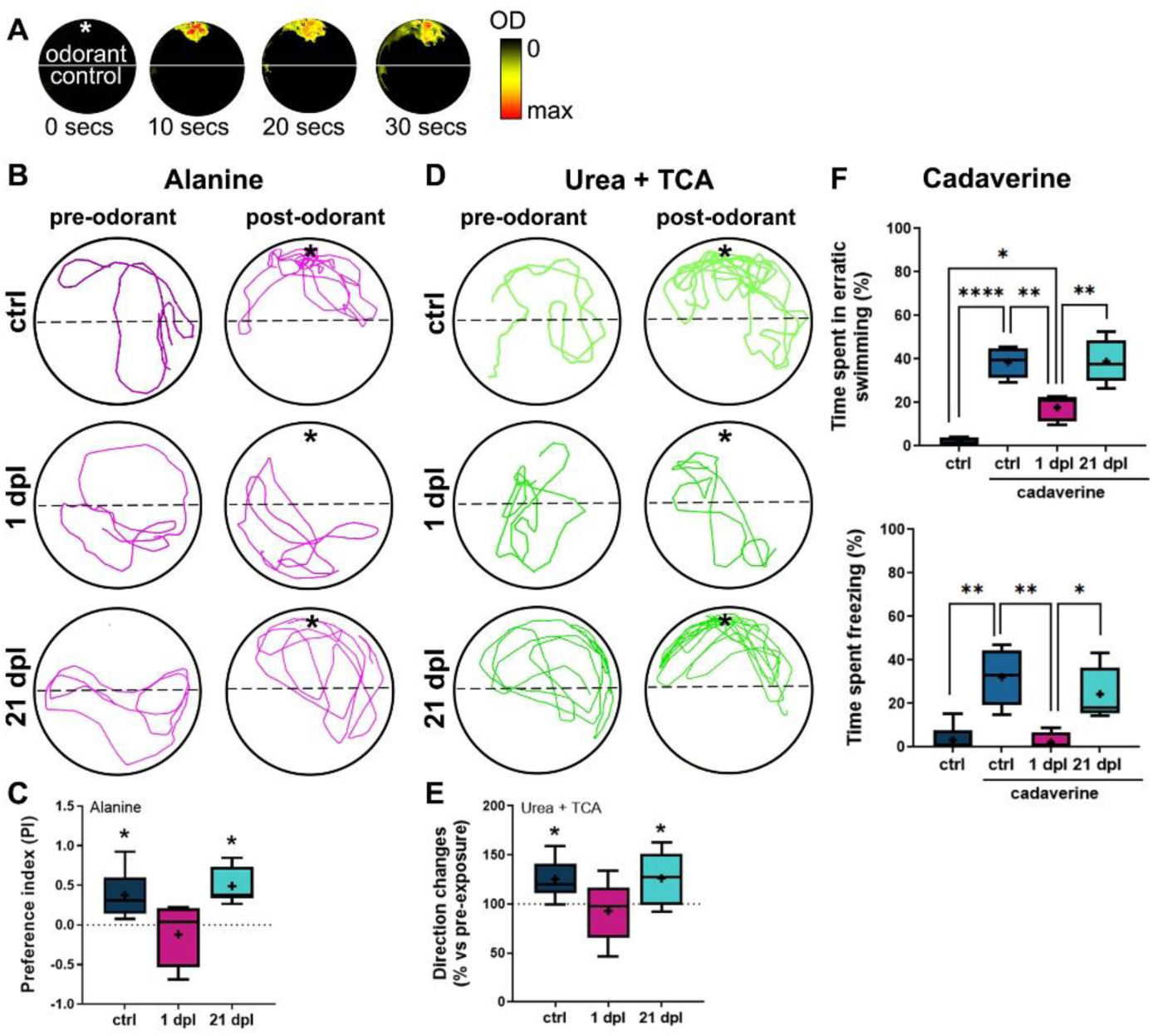
Effects of QA bulbar lesion on olfactory-mediated behavioral responses. A) Time course of a dye distribution in the behavioral chamber showing that odorant solutions remain in half of the chamber (indicated by an asterisk, *) during 30 seconds after delivery. In figures B) and C), the asterisk (*) indicates the position where the odorant solution was administered. B) Representative swimming trajectories of zebrafish pre-(30 secs) and post-(30 secs) alanine delivery in unlesioned controls (upper panels), 1-dpl (middle panels) and 21-dpl fish (lower panels). C) Quantification of preference index (PI) after alanine exposure compared against pre-odorant exposure from (B) (n = 5-6). One-sample Wilcoxon test, *p < 0.005 when compared to baseline of 100% (no change). D) Representative swimming trajectories of zebrafish pre-(30 secs) and post-(30 secs) exposure to a mixture of urea and taurocholic acid (TCA) in unlesioned controls (upper panels), 1-dpl (middle panels) and 21-dpl fish (lower panels). E) Quantification of percent change in swimming direction after exposure to a mixture of urea and TCA compared with pre-odorant exposure from (D) (n = 5-6). One-sample Wilcoxon test, *p < 0.005 when compared to baseline of 100% (no change). F) Quantification of responses to cadaverine in unlesioned controls, 1-dpl and 21-dpl fish. Top panel: percentage of time spent in erratic swimming after cadaverine exposure. Bottom panel: percentage of time spent freezing after cadaverine exposure. One-way ANOVA, *p < 0.005, **p < 0.001, ****p < 0.0001. Box plots indicate mean (+), quartiles (boxes) and range (whiskers).

In the 1-dpl group, olfactory-mediated behavioral responses to alanine and urea with TCA were impaired, indicating a loss of sensitivity to these odorants (Figs. 3 B-F). Interestingly, while the freezing aversive response to cadaverine was impaired, erratic swimming was only partially dampened, as it is significantly different from controls (Fig. F, top panel). This suggests that fish in the 1-dpl group retain some capacity to respond to cadaverine but not to the other odorants tested. Consistent with our prediction and our previous data on structural recovery, fish regained their olfactory function for all tested odorants by 21 dpl (Fig. 3 B-F). These results demonstrate that QA bulbar lesions impair olfactory function, which recovers along structural remodeling and recovery.

### QA lesions induce cellular proliferation and neurogenesis in the ventricular zone and the OB

The telencephalic ventricular zone (VZ) is known for its exceptional neurogenic capability, contributing to regenerative responses following damage in the telencephalon (Adolf et al., 2006; März et al., 2011). Similarly, the OE exhibits remarkable neurogenic abilities that mediate regeneration after injury or axotomy (Iqbal & Byrd-Jacobs, 2010; Kocagöz et al., 2022). Considering the extensive recovery and reorganization observed in our model (Figs. 1 and 2), we aimed to understand whether cell proliferation and migration contribute to post-lesion recovery. In zebrafish, newly born cells generated in the VZ migrate rostrally to the adjacent OB in an analogous route to the rodent rostral migratory stream (RMS) (Kishimoto et al., 2011). Thus, we hypothesized that OB lesions would stimulate proliferative responses in neural precursor cells (NPCs) within the VZ, differentiation of some of these cells into neurons, and migration to the OB. We also predicted that QA lesions would stimulate cell proliferation in the OE.

To investigate this, we assessed active proliferation using an antibody against PCNA (proliferating cell nuclear antigen) and tracked newly generated cells with the mitotic marker BrdU (bromodeoxyuridine). We examined the number of proliferative cells (PCNA+) in the entire telencephalon and within specific pallial and subpallial (i.e., ventrodorsal (Vd), and ventroventral (Vv)) regions (Fig. 4A). We observed a substantial increase in PCNA+ cells in the ipsilateral pallium of 1-dpl fish compared to the contralateral side and control fish (Fig. 4C, upper panel). Further analysis revealed increased proliferation in both subpallial regions in the lesioned hemisphere of 1-dpl fish (Fig. 4B, C middle and lower panels). Notably, the Vd region showed increased proliferation in both hemispheres (Fig. 4B, upper panels; C, middle panel). Overall telencephalic proliferation did not persist through 21 dpl, suggesting a homeostatic control of post-lesion cell proliferation (Fig. 4 B-D).

**Figure 4.**
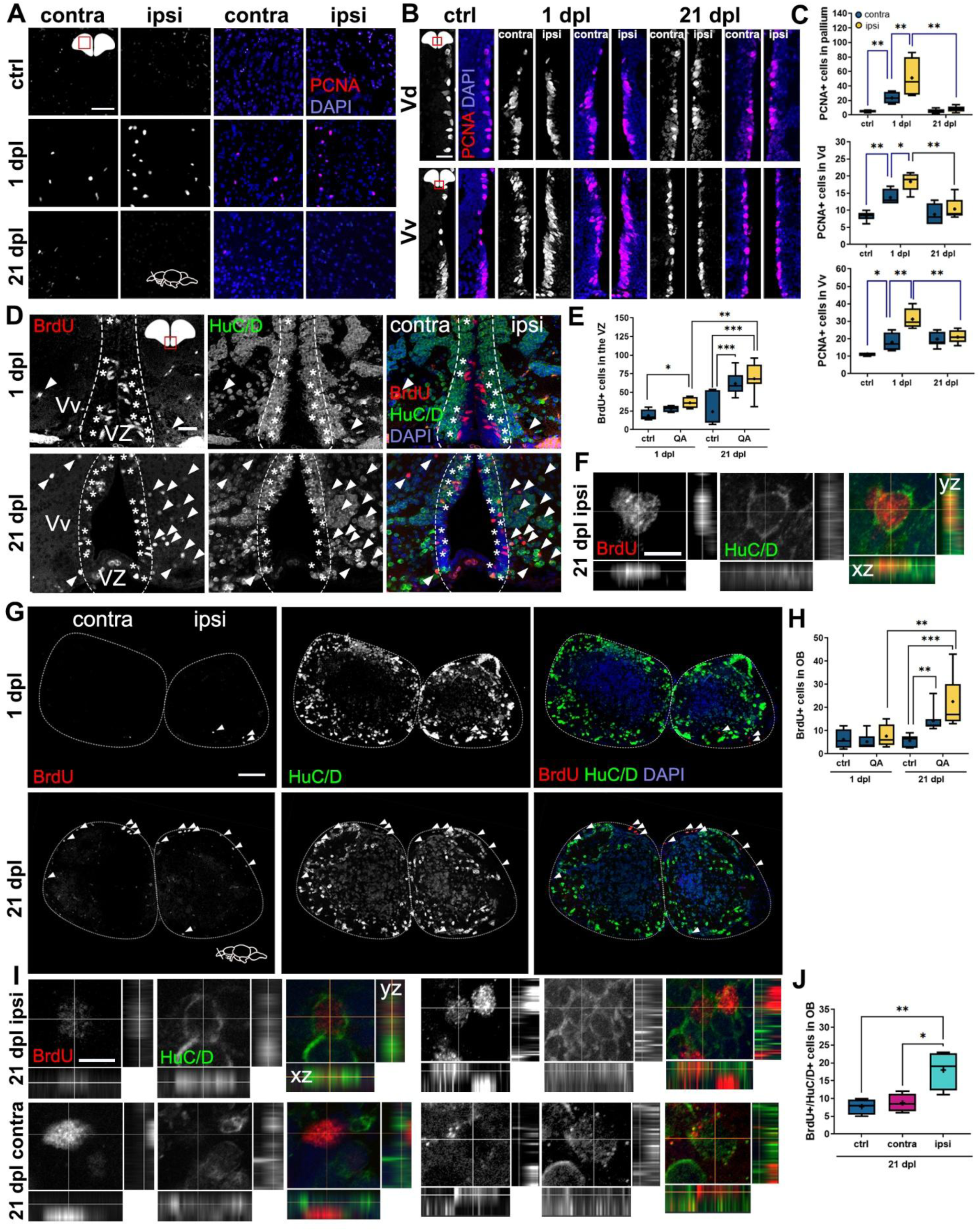
QA lesions to the olfactory bulb cause cellular proliferation and neurogenesis in the telencephalon and the OB. A) PCNA (proliferating cellular nuclear antigen) immunohistochemistry in coronal sections of the telencephalic pallium of unlesioned controls (top panels), 1-dpl (middle panels) and 21-dpl (lower panels) groups. Red: PCNA; Blue: DAPI. Scale bar: 100 µm. B) PCNA immunohistochemistry in coronal sections of the dorsal (Vd, top panels) and ventral (Vv, bottom panels) nuclei of the ventral telencephalon of unlesioned controls (left panel), 1-dpl (middle panels) and 21-dpl (right panels) groups. Red: PCNA; Blue: DAPI. Scale bar: 20 µm. C) Quantification of the number of PCNA+ cells in coronal sections of telencephalic regions from A) and B). Top panel: pallium, middle panel: Vd region, bottom panel: Vv region (n = 4-6). D) Double immunohistochemistry of BrdU and HuC/D in the neurogenic niche of the VZ (ventricular zone, indicated by dashed lines) and the Vv of 1-dpl (top panels) and 21-dpl (lower panels) groups. Cell migration indicated by arrowheads. BrdU+ progenitors in the VZ indicated by asterisks (*). Red: BrdU; Green: HuC/D; Blue: DAPI. Scale bars: 20 µm. E) Quantification of the number of BrdU+ cells in coronal sections of the VZ from (D) (n = 4-7). F) Orthogonal projection (xz, yz) of magnified views from the lower panel of (D) showing a migrating BrdU+/HuC/D+ newborn neuron in the ipsilateral telencephalon of a 21 dpl fish. Scale bar: 10 µm. G) Double immunohistochemistry of BrdU and HuC/D in the OB of 21-dpl group (lower panels). Red: BrdU; Green: HuC/D; Blue: DAPI. Scale bars: 20 µm. H) Quantification of the number of BrdU+ cells in coronal sections of the OB from (G) (n = 4-7). I) Orthogonal projections (xz, yz) of magnified views from (G) showing BrdU+/HuC/D+, BrdU+, and HuC/D+ cells in the contra- and ipsilateral OB of 21-dpl fish. Scale bar: 10 µm. J) Quantification of the number of BrdU+/HuC/D+ double-labeled cells in coronal sections of the OB from (G) (n = 4). Box plots indicate mean (+), quartiles (boxes) and range (whiskers). One-way ANOVA, *p < 0.005, **p < 0.001, ***p = 0.0009, ****p < 0.0001.

To study the fate and localization of post-lesion newborn cells at 1 dpl and 21 dpl, we employed a single BrdU pulse- and-chase assay to track cells generated (Fig. 4E), by administering BrdU intraperitoneally (i.p.) immediately after the QA lesion. Our BrdU exposure was limited to a single administration to avoid high fish mortality rates associated with multiple injections or BrdU immersion.

Additional control groups received BrdU without lesioning to discriminate between newly-generated cells due to the lesion specifically to those generated constitutively (Figs. S4A, B). We first examined BrdU+ cells in the neurogenic niche of the telencephalic ventricular zone (VZ) in lesioned fish and found a significant increase in newborn cells in the ipsilateral side at 1 dpl compared to the contralateral side and control fish (Figs. 4D, E). At 21 dpl, there was a significant increase in BrdU+ cells in both hemispheres compared to 1 dpl (Fig. 4H, right panel; G), with no difference between unlesioned controls at 1 dpl and 21 dpl (Fig. 4G). The approximately two-fold increase in BrdU+ cells from 1 dpl to 21 dpl suggests an additional round of cell division between 1 dpl and 21 dpl. This is supported by our PCNA+ findings, in which we observed robust proliferation ongoing at 1 dpl, likely after BrdU is no longer bioavailable (Figs. 4B, C; (Zupanc & Horschke, 1995).

To assess neurogenesis, we used the neural marker HuC/D along with BrdU. In addition to BrdU-labeled progenitors remaining in the VZ (Fig. 4D, asterisks), we observed BrdU+ cells migrating distally from the VZ in the ipsilateral hemisphere at 21 dpl (Fig. 4D, lower panel; arrowheads), supporting our hypothesis of cell migration. Some of these migrating cells were immunoreactive against both BrdU and HuC/D, indicating neurogenesis (Fig. 4D, lower panel), as confirmed by orthogonal projections in higher magnification images (Fig. 4F).

Next, we assessed BrdU+ profiles in the OB. There was no difference between lesioned and unlesioned groups at 1 dpl, or among OBs of unlesioned controls at 21 dpl (Figs. 4G, H). However, we found a significant increase in BrdU+ cells in both the contralateral and ipsilateral OBs at 21 dpl (Fig. 4H). The robust proliferation and generation of BrdU+ cells in the VZ, along with the migration of BrdU+ cells to the OB, support our prediction that most of these newborn cells originated in the VZ and migrated to the OB via the RMS (Kishimoto et al., 2011).

To assess neurogenesis in the OB, we counted BrdU+/HuC/D+ cells at 21 dpl and found evidence of neurogenesis in both control and lesioned groups, with a significant increase in the lesioned side (Figs. 4G, J). Higher magnification images revealed variation in BrdU signal intensity within nuclei, suggesting different amounts of BrdU incorporated into the cells’ DNA (Fig.4I). These results support our hypothesis that some BrdU+ cells labeled during the pulse- and-chase assay underwent an additional proliferation cycle. Overall, by 21 dpl, we found an increased number of newly born cells in the OB in both unlesioned controls and lesioned fish, with a significant increase of newborn neurons in the ipsilateral side (Fig. 4J), including some cells labeled with BrdU+/TH+ (Fig. S4C). These results indicate that neurogenesis is amplified by the bulbar lesion as a repair mechanism.

### QA lesions induce cellular proliferation and neurogenesis in the olfactory epithelium

Given the complete recovery of the OE by 21 dpl (Figs. 2C-E), we assessed cell proliferation and neurogenesis by tracking BrdU+ cells across different lamellar regions of the olfactory organ. We observed a significant increase of BrdU+ cells in the ipsilateral, but not the contralateral, lamellae at 1 dpl compared to unlesioned controls (Figs. 5A, D). The number of BrdU+ cells in unlesioned controls remained consistent between 1 dpl and 21 dpl (Fig. 5D). However, in the contralateral side of 21 dpl fish, the number of BrdU+ cells decreased compared to both the 1 dpl group and the ipsilateral side at 21 dpl (Fig. 5D). Importantly, there was no significant difference in the number of BrdU+ cells in the ipsilateral OE between 1 dpl and 21 dpl (Fig. 5D). These results indicate that while BrdU+ cell numbers in the ipsilateral OE remain constant from 1 dpl to 21 dpl, the number of BrdU+ cells on the contralateral side decreases over time. This implies that cell proliferation is robust in both lamellae but that the excess of newborn cells generated on the contralateral side are likely eliminated as they are not needed to replace lost neurons, consistent with previous results (Bayramli et al., 2017; Kocagöz et al., 2022).

**Figure 5.**
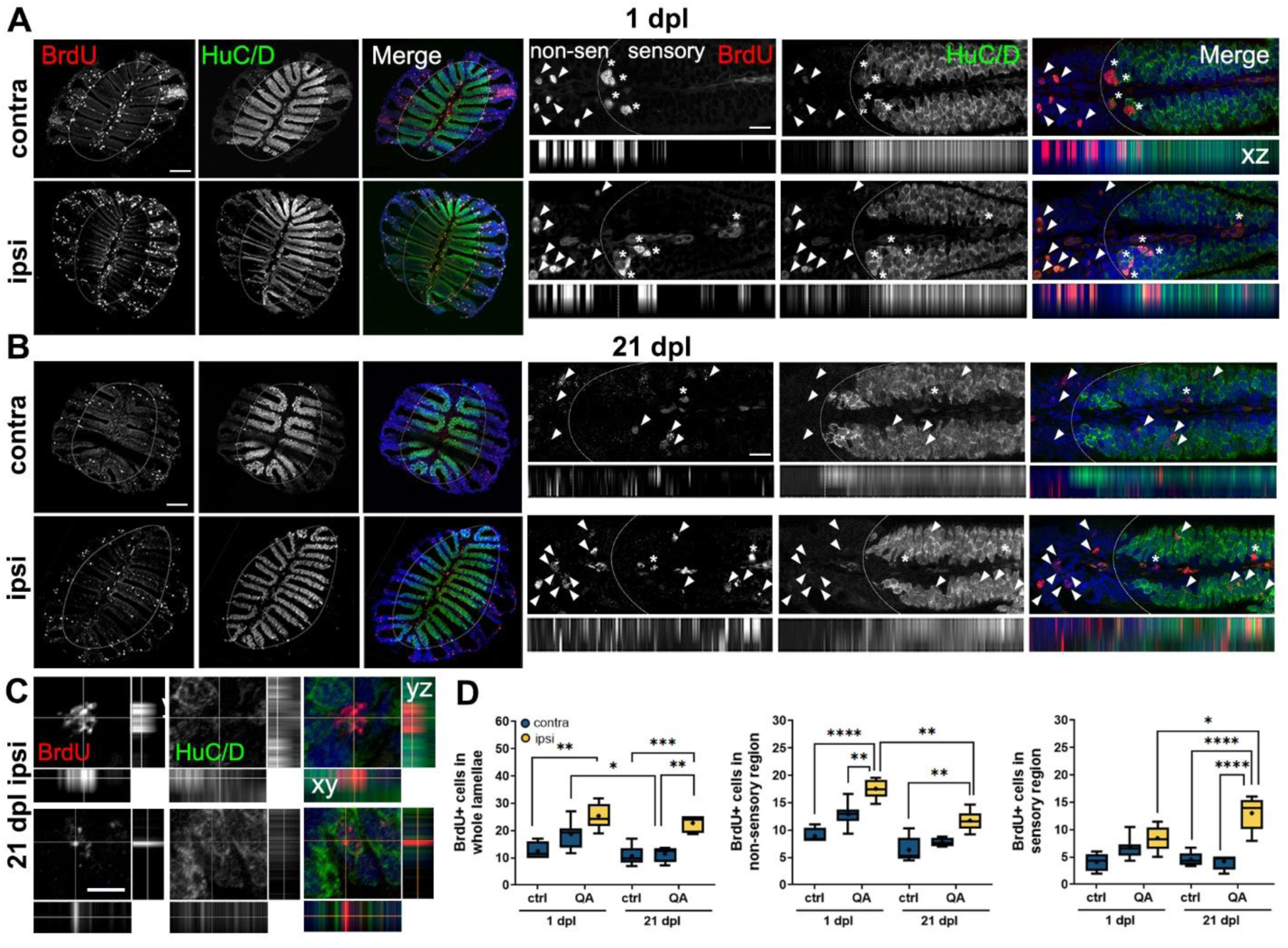
QA lesions to the olfactory bulb cause cellular proliferation and neurogenesis in the OE. A) and B) Left panels: Double immunohistochemistry of BrdU and HuC/D in sections of the whole olfactory organ of 1-dpl (A) and 21-dpl (B) groups. Right panels: Magnified views of left panels, showing the sensory and non-sensory regions of the OE lamellae (indicated by dashed lines). Below each panel, orthogonal (xz) projections are shown. BrdU+ progenitors in the sensory area are indicated with asterisks (*). Red: BrdU; Green: HuC/D; Blue: DAPI. Scale bar: 100 µm. C) Orthogonal projections (xz, yz) of magnified views from B) showing BrdU+/HuC/D newborn neurons in the ipsilateral sensory region of 21-dpl fish. Scale bar: 10 µm. D) Quantification of the number of BrdU+ cells in OE sections from (A) and (B). Left panel: whole lamellae, middle panel: non-sensory region, right panel: sensory region (n = 4-7). Box plots indicate mean (+), quartiles (boxes) and range (whiskers). One-way ANOVA, *p < 0.005, **p < 0.001, ***p = 0.0009, ****p < 0.0001.

To assess neurogenesis in the OE, we examined the location of post-lesion newly generated cells within the epithelium. Newborn cells that acquire a neuronal fate in the OE migrate first horizontally to the sensory region, then apically as they integrate into the sensory epithelium (Kocagöz et al., 2022). We quantified BrdU+ cells in the non-sensory and sensory regions of the lamellae that are demarcated by the absence and presence of the neuronal marker HuC/D, respectively (Fig 5A, indicated by dotted white lines in whole olfactory organs and individual lamellae). In the ipsilateral OE at 1 dpl, there was a significant increase in BrdU+ cells in the non-sensory region, where globose basal cells (GBCs) are located (Figs. 5A, D;(Kocagöz et al., 2022). In contrast, no significant difference was observed in the sensory region of the ipsilateral side (Figs. 5A, D). Notably, we observed large BrdU+ cells in the sensory/non-sensory border of the lamellae, appearing to be actively dividing, based on their nuclear morphology (Fig. 5A, asterisks). In addition, we observed BrdU+ cells in the basal laminae of the ipsilateral OE (Fig. 5A, middle panels, asterisks), suggesting that these cells are HBCs (horizontal basal cells) activated by epithelial damage (Kocagöz et al., 2022).

At 21 dpl, there was a decrease in BrdU+ cells in the ipsilateral non-sensory lamellae compared to the same region at 1 dpl (Figs. 5A, B, D middle panel), though the number of BrdU+ cells at 21 dpl remained significantly different from controls (Fig. 5D, middle panel). Conversely, there was a significant increase of BrdU+ cells in the ipsilateral sensory epithelium at 21 dpl (Fig. 5B, arrowheads; D right panel). Some of these BrdU+ cells were localized towards the apical region of the epithelium. Orthogonal image projections confirmed that BrdU+ cells that migrated horizontally and apically in the sensory region overlapped with HuC/D staining (Fig. 5C), indicating integration of these newborn neurons into the sensory OE.

Additionally, similar to our observations in the OB, we noted variations of the intensity of BrdU signal in cells in both sensory and non-sensory regions of the epithelium at 21 dpl. While GBCs in the non-sensory region and in the sensory/non-sensory border exhibited intense BrdU signal in the nucleus at 1 dpl (Fig.5A, arrowheads and asterisks), the intensity of the signal faded by 21 dpl in some cells (Fig. 5B, arrowheads). BrdU+/HuC/D+ cells in the sensory epithelium also showed different levels of signal intensity, often appearing as discrete nuclear puncta (Fig. 5C). These observations suggest that BrdU-labeled progenitors have undergone additional cell divisions between 1 dpl and 21 dpl.

### QA lesions induce neuroinflammation in the olfactory system and telencephalon, which mostly subsides within 21 days

We investigated neuroinflammatory responses in the OB, OE, and telencephalon, as neuroinflammatory modulators are known to underlie some regenerative and repair processes following injury in brain regions such as the telencephalon (Kyritsis et al., 2012; Marz et al., 2010). Astroglial and microglial responses in the OB following damage to the OE have also been characterized (Scheib & Byrd-Jacobs, 2020; Var & Byrd-Jacobs, 2019). We hypothesized that QA lesions of the OB would induce neuroinflammatory responses potentially associated with repair responses observed.

To assess astroglial activation, we examined the glial marker GFAP (glial fibrillary acidic protein). In the zebrafish OB, GFAP is expressed in glial cells that lack the stellate morphology seen in mammalian brains. Instead, they are characterized by diffuse, thin processes throughout the OB, with higher staining in the outermost olfactory nerve layer of the OB (Lazzari et al., 2014; Scheib & Byrd-Jacobs, 2020). At 1 dpl, we observed induction of astroglial activation (indicated by increased anti-GFAP optical density, OD) in the lesioned side compared to unlesioned controls, with the activation returning to pre-lesioned levels by 21 dpl (Figs. 6A, B). Similarly, we observed astroglial activation in the telencephalon at 1 dpl, subsiding by 21 dpl (Figs. 6C, D). No GFAP staining was observed in the OE (data not shown), consistent with reports of astroglial cells being absent in the peripheral epithelium (Lazzari et al., 2014).

**Figure 6.**
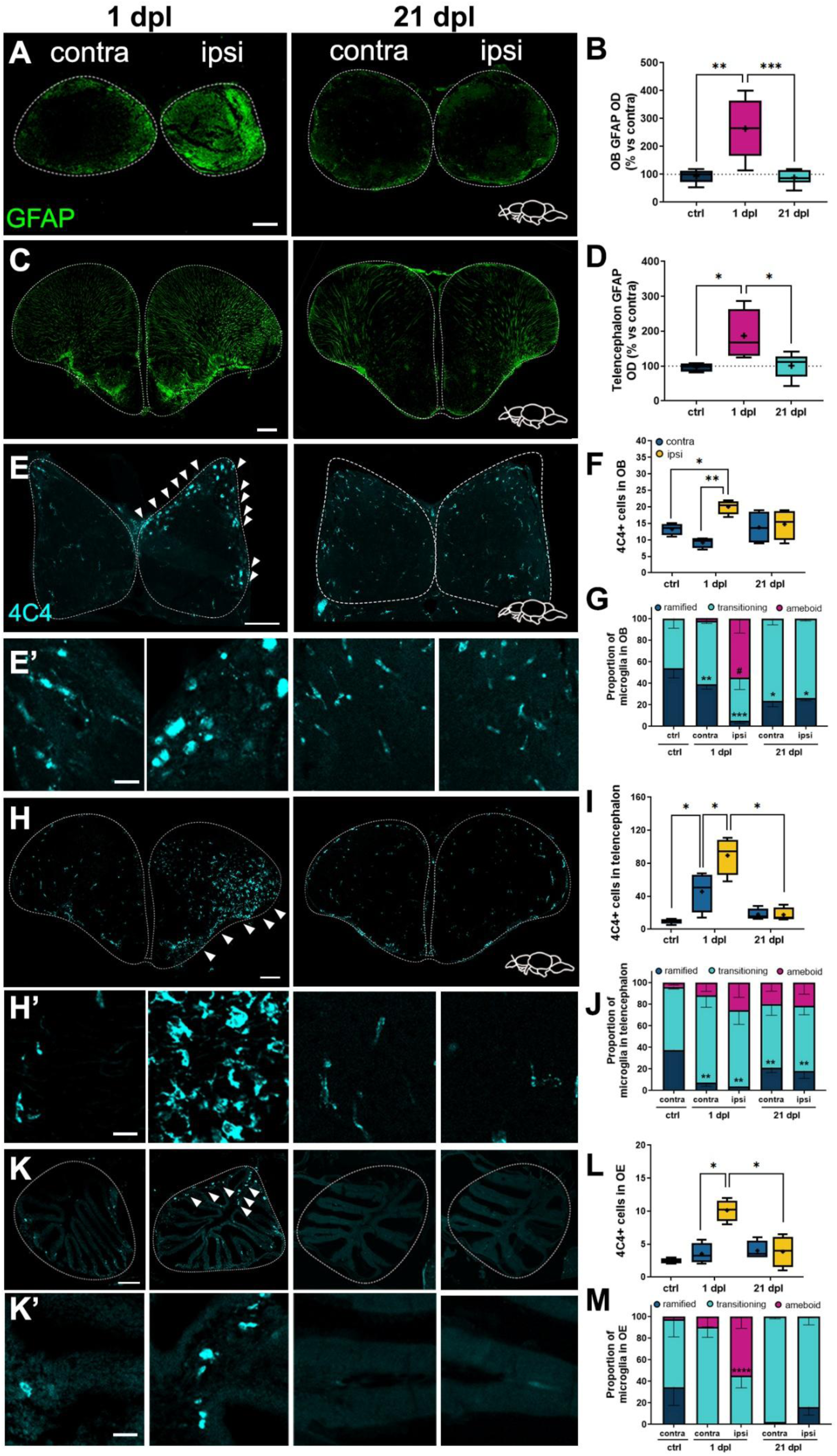
QA lesion leads to neuroinflammatory responses in the OB, telencephalon, and OE. A) and C) GFAP (glial fibrillary acidic protein) immunohistochemistry in coronal sections of the OB (A) and the telencephalon (C). Green: GFAP. Scale bar: 100 µm. B) and D) Quantification of percent change (vs. contra) in optical density (OD) of GFAP in the OB (B) and telencephalon (D) (n = 5-7). E), H) and K) 4C4 immunohistochemistry in horizontal sections of the OB (E), coronal sections of the telencephalon (H), and sections of the OE (K). Cyan: 4C4. Scale bar: 100 µm. E’), H’) and K’) Magnified views of the corresponding panels (E, H, K) indicate microglial morphology. Cyan: 4C4. Scale bars: 20 µm. F), I) and L) Quantification of average 4C4+ cells in the OB (F), telencephalon (I) and OE (L) (n = 4). Box plots indicate mean (+), quartiles (boxes) and range (whiskers). One-way ANOVA, *p < 0.005, **p < 0.001, ***p = 0.0009, ****p < 0.0001. G), J) and M) Quantification of the proportion (out of 100%) of microglial morphologies in the OB (G), telencephalon (J) and OE (M) (n = 4). Values represent means ± SEM. One-way ANOVA. In (G) *p < 0.005, **p < 0.001, ***p = 0.0009 in ramified microglia in 1-dpl and 21-dpl groups against unlesioned controls. #p < 0.0001 in activated microglia in the ipsilateral OB of 1-dpl group compared to activated microglia from all groups, ****p < 0.0001 in the activated microglia in the ipsilateral OB of 1-dpl group compared with activated microglia from all groups. In (J), **p < 0.001 in ramified microglia in 1-dpl and 21-dpl groups against unlesioned controls. In (M), ****p < 0.0001 against ameboid microglia compared with all other groups.

Next, we assessed microglial activation and migration in the OB, OE, and telencephalon by using the monoclonal antibody 4C4 that recognizes the microglia marker, galectin 3-binding protein (Rovira et al., 2023). At 1 dpl, we found a substantial increase in 4C4+ cells in all regions studied (Figs. 6E, F, G, H, I, J). Interestingly, we observed a significant increase in microglial cells in the contralateral hemisphere of the telencephalon at 1 dpl (Figs. 6G, H), although we did not observe the same trend in the OE or in the OB (Figs.6E, F, I, J). By 21 dpl, the microglial response had subsided in all regions (Figs. 6E, F, G, H, I, J). Microglial responses are characterized by both the number of cells present at the site of damage and their activation state (Var & Byrd-Jacobs, 2019). Resting microglia exhibit a ramified morphology, characterized by a small soma and thin, ramified processes. In contrast, activated microglia transform into phagocytic ameboid cells with a large soma and minimal to no processes. A transitioning morphology between these states is marked by an enlarged soma and shorter, fewer processes (Jonas et al., 2012; Var & Byrd-Jacobs, 2019). Using this classification, we assessed and quantified microglial morphologies across the OB, OE, and telencephalon. In the OB, we noted a significant increase in activated amoeboid cells on the ipsilateral side at 1 dpl, along with a rise in transitioning profiles in the contra- and ipsilateral sides in the contralateral side at 21 dpl and both hemispheres at 21 dpl (Figs. 6E, E’, F). Most amoeboid 4C4+ cells were found along the olfactory nerve layer on horizontal sections of the OB (Fig. 6E arrowheads), indicative of ipsilateral migration through olfactory axons. In the telencephalon, there was a decrease in ramified, resting microglia in both hemispheres at 1 dpl and 21 dpl compared to controls (Figs. 6G, G’, H). In the OE, we found a significant shift into an activated amoeboid morphology in the ipsilateral OE at 1 dpl, with morphology returning to normal by 21 dpl (Figs. 6I, I’, J).

## DISCUSSION

In this study, we employed a novel lesioning method of the olfactory bulb by unilateral injection of the endogenous excitotoxic agent quinolinic acid (QA). This approach allowed us to investigate mechanisms of olfactory degeneration and regeneration in adult zebrafish.

Two major findings emerge from this work. First, our model effectively reproduces key features of brain-injury-induced olfactory dysfunction. We observed significant degeneration and neuronal loss in the lesioned OB, resulting in retrograde degeneration and loss of OSNs in the OE. This widespread degeneration leads to a loss of olfactory function.

Second, we report that by 21 dpl, the olfactory system displays outstanding structural reorganization, regeneration, and recovery in the OB and OE, concomitant with complete restoration of olfactory function. This recovery is supported by substantial regenerative and plasticity mechanisms in the olfactory system, including neurogenesis.

### Effects of the QA lesion on olfactory bulb’s morphology and neurons

In our model, QA lesions primarily target glutamatergic bulbar mitral cells, as expected, given their susceptibility to excitotoxic damage (Li et al., 2022; Schubert et al., 2022). Mitral cells play a crucial role in olfactory perception, as they receive and integrate inputs from OSNs and relay this information to higher olfactory centers for further processing (Miyasaka et al., 2014). An interesting finding is that although Tbr2a+ mitral cells on the lesioned side at 21 dpl do not fully recover to control levels, there is a significant difference in cell number between 1 dpl and 21 dpl groups. Regeneration of mitral cells in adult organisms has not been described before; however, our finding raises the possibility that mitral cells may regenerate post-lesion in adult zebrafish. Given that we used a single injection of BrdU, the number of BrdU+ newly born cells generated post-lesion is limited; this might explain why we did not find any BrdU+/Tbr2a+ cells. Further investigations using alternative experimental strategies are required to elucidate whether mitral cells can recover in this lesion paradigm.

Our results also indicate that dopaminergic periglomerular neurons are resistant to the excitotoxic lesion, aligning with previous observations of catecholaminergic cells’ resistance to QA-induced neurotoxicity (Mazzari et al., 1986; Venero et al., 1995). We observed, however, that the QA lesion considerably alters the localization of Tbr2a+ somata and periglomerular cells within bulbar layers, due to the significant reduction in bulb volume and neuropil degeneration. By 21 dpl, the bulbar volume, laminar organization, and appearance, return to control levels.

### Effects of the QA lesion on olfactory glomeruli

We hypothesized that the loss of mitral cells would impact the structure of olfactory glomeruli, as the adequate localization of dendritic puncta is critical for effective synaptic contacts with axonal termini, a phenomenon observed in other models of excitotoxicity (Lundberg et al., 1994). Indeed, we found that the lesion caused significant disorganization and defasciculation of dorsal and dorsolateral olfactory glomeruli, while ventral glomerular clusters remained unaffected. By 21 dpl, the overall organization and morphology of the dorsolateral glomeruli had recovered. However, Gαolf-positive glomeruli in the dorsal cluster still remained slightly disorganized and defasciculated. These findings align with previous reports indicating that chemical lesioning of the OE disrupts olfactory glomeruli in the OB, with most glomeruli recovering except those in the dorsal and medial clusters (White et al., 2015). The dorsal glomerular cluster conveys sensory information from ciliated OSNs, responsive to bile salts (Sato et al., 2005; Braubach et al., 2012; Masuda et al., 2024). Interestingly, these OSNs have been reported to be inherently susceptible to various types of chemical and metal-induced damage to the OE (White et al., 2005; Hentig and Byrd-Jacobs, 2016; reviewed in Calvo-Ochoa and Byrd-Jacobs, 2019). This phenomenon has also been observed in mammals, where OE degeneration resulting from axotomy leads to differential timing of glomerular degeneration (Graziadei & Monti Graziadei, 1980). The reason for the differential effects of the lesion on specific glomerular clusters remains to be elucidated.

### Effects of the QA lesion on the olfactory epithelium

After characterizing the lesion-induced degeneration in the OB, we hypothesized that OB output cell loss and OSN axonal damage would retrogradely lead to OSN degeneration, as reported in models of olfactory axotomy and bulbectomy (Graziadei and Graziadei, 1979; Schwob et al., 1992; Kocagoz et al., 2022). As expected, we observed substantial degeneration in the sensory region of the OE, along with notable disruption of OSN morphology and organization within the epithelial layers. Consistent with our findings in the OB, the OE appeared fully recovered by 21 days, with OSNs returning to pre-lesioned morphology and organization.

### Effects of the QA lesion on olfactory function

We confirmed that olfactory function is impaired due to the extensive neurodegeneration observed across the olfactory system following the QA lesion. We assessed olfactory responses to three odorants with physiological relevance for fish, processed by two major types of OSNs and different glomerular clusters: alanine, urea and taurocholic acid (TCA), and cadaverine. Interestingly, lesioned fish showed a loss of olfactory-mediated responses to alanine, urea and TCA, and a partial loss of response to cadaverine. This indicates that unilateral degeneration caused by the QA lesion is sufficient to result in impairment of olfactory function. Previous studies have also shown loss of olfactory responses following unilateral OE exposure to toxicants in fish (Kolmakov et al., 2009; Abreu et al., 2017).

Olfactory function is restored within 21 days, indicating that the olfactory system recovers both structurally and functionally. Although Tbr2a+ mitral cell numbers and glomeruli did not completely return to pre-lesioned levels by this time, the olfactory system appears capable of remodeling and reorganizing to support functional recovery, highlighting its remarkable levels of neuroplasticity (White et al., 2015). In line with this, zebrafish mitral cell dendritic arbors have shown to be repaired and restored following processes of injury-induced deafferentation that leads to a decrease of their distribution, arborization, and length (Pozzuto et al., 2019; Rozofsky et al., 2024). Notably, after 8 weeks post-injury, mitral cell dendritic arbors are larger than those of controls, independent of animal growth (Rozofsky et al., 2024), underscoring the resilience and neuroplastic abilities of these cells.

### Effects of QA lesion on proliferation and neurogenesis

This study is the first to demonstrate an integrated proliferative and neurogenic response across the telencephalon, OB, and OE, underlying the structural and functional recovery of the olfactory system after brain injury. This adds to the current literature, which has described in situ proliferative mechanisms in these regions individually: the telencephalon (Kroehne et al., 2011; Skaggs et al., 2014), the OB (Villanueva and Byrd-Jacobs 2009; Trimpe and Byrd-Jacobs 2016), and the OE (Bayramli et al. 2017; Kocagoz et al., 2022). Previous reports have established that repair of injured neural tissue requires inflammatory responses, progenitor proliferation and neurogenesis, and migration of newborn neurons to the lesioned areas (reviewed in Calvo-Ochoa et al., 2020), all of which were observed in our model.

While the proliferative response was robust in both hemispheres of the VZ of the telencephalon, the OB, and the OE, we found increased neurogenesis in the lesioned side by 21 dpl. This indicates neuronal replacement and integration into the repairing tissue, consistent with previous findings (Kroehne et al., 2011; Skaggs et al., 2014; Trimpe and Byrd-Jacobs 2016). An important consideration, as mentioned in the results section, is that our BrdU pulse-and-chase assay was limited to a single BrdU administration. Consequently, our results provide an important, but limited, view of newly born cells and neurons generated after the QA lesion. Alternative methods to label newly generated cells for longer periods of time will yield a more complete picture about the extent of neurogenesis and neural integration in the recovered olfactory system.

An important conclusion from our data is that the number of newborn cells observed in the OB and OE of 21-dpl groups indicate further division of post-lesioned BrdU+ cells after 1 dpl. This observation is supported by studies employing continuous or longer BrdU exposure to track cell proliferation across several hours or days, showing that both constitutive and post-injury proliferation in the brain and OE are sustained (Zupanc et al., 2005: Hinsch and Zupanc, 2007; Kroehne et al., 2011; Bayramli et al. 2017; Kocagoz et al., 2022). In line with this, it has been reported that progenitors in the VZ of Vv and Vd regions (type IIIa and IIIb, respectively) can undergo at least two rounds of cell division followed by a final symmetric division that gives rise to neuroblasts migrating to the OB (Rothenaigner et al. 2011; Kishimoto et al., 2011).

### Effects of QA lesion on neuroinflammation

Neuroinflammation serves as a critical modulator of proliferation and repair in the zebrafish brain (Kyritsis et al., 2012; Tsarouchas et al., 2018). Here, we report that QA lesions induce neuroinflammatory responses through astroglial and microglial activation in the OB, OE, and telencephalon. GFAP+ radial glia-like cells in the outer ventricular surfaces of the telencephalon, including the VZ, are bona fide neural precursor cells NPCs (Zupanc and Clint, 2003; Marz et al., 2010; Rothenaigner et al., 2011; reviewed in Calvo-Ochoa et al., 2020). In models of telencephalic injury, a robust astroglial response has been reported in the telencephalon, along with cell proliferation in the VZ (Zupanc and Clint, 2003; Marz et al., 2010; Skaggs et al., 2014). This is consistent with our findings, suggesting that injury in the OB induces a similar neuroimmune-proliferative response to the one reported in the lesioned telencephalon. Future investigation is needed to elucidate whether the GFAP+ astroglia activated after lesions in the OB also modulate regenerative responses in the olfactory system.

Additionally, we also observed enhanced activation and recruitment of microglia in the ipsilateral hemisphere and OE at 1 dpl, with a dampened but sustained response in the OB and telencephalon by 21 dpl. At 1 dpl, the ipsilateral OB experienced a dramatic recruitment of activated, amoeboid microglia, indicative of phagocytosis (Herbomel et al., 1999; Peri and Nüsslein-Volhard, 2008). These amoeboid microglia were found along the peripheral layer of the bulb, suggesting rostral migration towards the site of injury via olfactory axons. By 21 dpl, we observed remodeling of olfactory glomeruli and neurogenesis in the OE, prompting us to propose that new OSNs extend projections to the OB and form new glomeruli (White et al., 2015). For this process to occur, degenerated axons and post-lesion debris must first be cleared. We posit that phagocytic microglia play a crucial role in this process, as they facilitate the establishment of new axonal connections by clearing out degenerated axonal termini in the zebrafish CNS (Stuermer et al., 1992; David and Kroner, 2011). In the telencephalon, we found an increased number of transitioning microglia between active and resting states and activated microglia in the lateral surface of the ipsilateral hemisphere at 1 dpl, suggesting recruitment of microglia from posterior brain areas, as previously described (Var and Byrd-Jacobs, 2019). Furthermore, some transitioning microglial profiles remained in both telencephalic hemispheres at 21 dpl, indicating a continued, active role of these neuroinflammatory cells in the maintenance of the recovering brain.

In conclusion, this study provides an integrated characterization of structural repair, regeneration processes, and functional recovery in the olfactory system following an excitotoxic lesion of the OB. We demonstrate that proliferation and neurogenesis in the OB, OE, and telencephalon support the recovery of the degenerated olfactory system and likely underlie the restoration of olfactory function. Our findings also support the hypothesis that excitotoxic damage in the OB serves as a mechanistic link between brain injury, neuronal loss in the OB, and olfactory dysfunction in rodent models (Marin et al., 2015 and 2022). Our work adds to the body of literature validating the use of zebrafish as a tractable model for investigating fundamental processes of neural plasticity, repair, and recovery in the olfactory system. Understanding the mechanisms underlying this prolific regenerative response could pave the way for innovative therapeutic approaches to promote olfactory recovery in patients experiencing olfactory dysfunction due to brain injuries.

## ACKNOWLEDGEMENTS

We are grateful to current and past members of the Calvo lab for support throughout the completion of these studies; in particular to Mackenzie Williams, Ashely Trainor, Matthew Czmer, and Carmen Casper for technical support. We acknowledge the following institutions and agencies that have generously funded this work. ECO was supported by: National Science Foundation (grant PRFB 1811447), International Brain Research Organization (IBRO, Rising Stars Award), IBRO-RIKEN CBS fellowship, and Hope College. NWV received an Anderson research fellowship from Hope College. SLD received a fellowship from the Grace A. Dow foundation. ABG received a Sherman Fairchild fellowship from Kalamazoo College. YY was supported in part by Grant-in-Aid for Scientific Research (20300117) from the Ministry of Education, Culture, Sports, Science and Technology of Japan. CBJ was supported by Western Michigan University.

## Author contributions

E.C.O. conceived the project, designed experiments, and performed QA lesions. E.C.O., N.W.V., and T.P.L performed tissue preparation, sectioning, antibody stainings, confocal microscopy, and analyzed imaging data. S.L.D., E.A.T., and A.B.G. performed behavioral assays and analyzed behavioral data. N.M., Y.Y., and C.B.J. offered expertise to various components of the project and provided reagents and equipment. E.C.O., N.W.V., N.M., Y.Y., and C.B.J contributed to writing and editing the manuscript. E.C.O., Y.Y., and C.B.J. acquired funding.

## MATERIALS AND METHODS

### Animals

Adult wild-type zebrafish (Danio rerio) of both sexes were kept and bred in a filtered aquarium system kept at 28° C (Aquaneering, San Diego, CA) located at Hope College’s zebrafish facility and in aquaria at Western Michigan University and RIKEN Center for Brain Science (CBS). The aquarium room was kept on a 12-hr light: 12-hr dark cycle. Fish were fed 3 times a day ad libitum, twice with commercial flake food (Aquaneering, San Diego, CA) and once daily with freshly hatched brine shrimp (Brine Shrimp Direct, Ogden UT). All experiments were carried out in accordance with the National Institutes of Health Guide for the Care and Use of Laboratory Animals (NIH Publication No. 80-23) and were approved by the Institutional Animal Care and Use Committees from Hope College, Western Michigan University, and RIKEN CBS. All efforts were made to minimize the number of animals used.

#### Olfactory bulb excitotoxic lesion

To generate focal unilateral excitotoxic damage to the olfactory bulb, adult fish were anesthetized by submersion in a 0.03% MS222 (tricaine) solution (Sigma-Aldrich, St. Louis, MO) until opercular movements slowed down and the fish were irresponsive to a tail pinch. Anesthetized fish were then subjected to a focal excitotoxic lesion by injection of 1 μl of 15 mM quinolinic acid (QA; Sigma-Aldrich, St. Louis, MO) with the use of a beveled syringe with at 30-gauge needle (Hamilton, Reno, NV), inserted diagonally through the skull into the right olfactory bulb (modified from Skaggs et al., 2014). Eye and nose position as well as visible skull sutures were used as guides for locating the olfactory bulbs. The left, non-lesioned bulbs served as internal controls. Fish were left to recover for either 1 or 21 days and then were euthanized by over-anesthetization with a 0.03% MS222 solution and decapitated, and heads were fixed with 4% paraformaldehyde (PFA; Sigma-Aldrich, St. Louis, MO) in PBS for 24 hours at 4°C. The next day, brains, olfactory epithelia or whole heads were dissected and treated for immunohistochemistry or whole mount preparations.

### Tissue processing

#### Sectioned tissue for immunohistochemistry

Dissected brains and olfactory organs were post-fixed in 4% PFA for 1 to 2 hrs. Then, they were subjected to 15-min incubations with solutions of increasing ethanol concentrations to slowly dehydrate the tissue, followed by an incubation in xylene (Sigma-Aldrich, St. Louis, MO). Dehydrated brains and olfactory organs were embedded in paraplast plus paraffin (McCormick Scientific, Berkeley, CA) and cooled to solidification. Then, semi-serial, 10-μm sections were obtained and adhered to charged slides (ThermoFisher, Waltham, MA). Brains were sectioned in the coronal plane while dissected olfactory organs were sectioned horizontally.

#### Whole head preparations

Fixed whole heads were dissected and decalcified with RDO rapid decalcifying solution (Electron Microscopy Sciences, Hatfield, PA) for 2 hrs, dehydrated through ascending ethanol washes, and embedded in paraffin, as described above. Semi-serial, 10-μm sections in the horizontal plane were obtained and then mounted on positively charged slides.

### Cell proliferation assays

To determine cell proliferation, we performed a pulse-and-chase assay with the thymidine analog, 5-bromo-2-deoxyuridine (BrdU, Sigma Aldrich). We administered intraperitoneally a single injection of a 10 μL of a 50 mg/ml BrdU solution in PBS immediately after the bulbar lesion. Fish in control groups received a BrdU injection but no bulbar lesions. After injections, fish were returned to fish water to recover.

### Immunohistochemistry

#### Sectioned tissue preparations

Mounted tissue was rehydrated by descending ethanol incubations and then subjected to antigen retrieval with a 10 mM sodium citrate (Sigma-Aldrich, Canada) solution (pH = 6.0) at 100°C for 10 min. Next, slides were washed with PBS and incubated for at least an hour with blocking buffer containing 3% normal goat serum (NGS; Vector Laboratories, Burlingame, CA) and 0.4% Triton X-100 (Sigma Aldrich, St. Louis, MO). Following blocking, slides were incubated with primary antibodies (Table 1) overnight and then incubated with fluorescently-labeled secondary antibodies for up to 2 hrs (Table 1). 4’,6-diamidino-2-phenylindole (DAPI; BD Pharmingen, Franklin Lakes, NJ) was used as a nuclear counterstain, added to the secondary antibody solution at 1 µg/ml. The sections were washed and then coverslipped using PVA-DABCO (Sigma Aldrich, St. Louis, MO), and examined with a confocal laser-scanning microscope Nikon A1 and the NIS-Elements software (Nikon, Japan).

**Table 1.** Antibodies used in this study.

| Antibody | Species | Dilution | Source |
| --- | --- | --- | --- |
| HuC/D | Mouse | 1:100 | ThermoFisher |
| Tbr2a | Rabbit | 1:500 | Yoshihara lab, RIKEN CBS |
| 4C4 | Mouse | 1:100 | Gift from Dr. Peter Hitchcock, University of Michigan |
| GFAP | Rabbit |  | DAKO |
| TH | Rabbit | 1:500 | Millipore |
| BrdU | Rat | 1:100 | Abcam |
| PCNA | Mouse |  | Sigma |
| Gaolf | Rabbit | 1:1000 | Yoshihara lab, RIKEN CBS |
| Calretinin | Goat | 1:1000 | Swant |
| SV2 | Mouse | 1:1000 | DSHB |
| Secondaries anti-mouse, rat, and rabbit IgG for immunohistochemistry | Goat | 1:200 | Invitrogen |
| Secondaries anti-mouse, goat, and rabbit IgG for whole-mounts | Donkey | 1:300 | Life technologies, Jackson laboratories, Biotium |

#### Whole mount preparations

Dissected and fixed brains were permeabilized by increasing methanol washes, subjected to antigen retrieval as described above, and incubated in blocking solution containing 5% normal horse serum (Antibodies Incorporated, Davis, CA) and 0.1% Triton X-100 (Sigma Aldrich, St. Louis, MO) overnight at 4°C. Then, the tissue was incubated with primary antibodies overnight (Table 1), washed, and incubated with fluorescently labeled secondary antibodies for 30 min at room temperature (RT) (Table 1). Following incubation, brains were kept away from light at 4°C in PBS-0.04% Triton X. Before confocal visualization, brains were mounted in 1.5% low-melting point agarose gel on glass coverslips. Whole olfactory bulbs were scanned every 3 m from dorsal and ventral sides using an inverted microscope (Olympus IX81) equipped with FV1000 confocal laser-scanning system. Imaging parameters used were: x10 dry objective lens (NA 0.4, Olympus UPlanApo), 512 x 512 pixels with x1.0 zoom.

### Histological analyses

For morphological analysis, sectioned tissue samples were deparaffinized with xylene, stained with Hematoxylin-Eosin (H&E; Vector laboratories, Newark, CA), and mounted with Permount (ThermoFisher, Waltham, MA) as previously described (Hentig and Byrd-Jacobs, 2016). These sections were examined under light microscopy with a Leica DM5000 B Automated Upright Microscope and an Application Suite X System software (Wetzlar, Germany).

### Olfactory bulb (OB) stereology

OB volumes were calculated using coronal sections stained with H&E by using a stereological Cavalieri estimator method (Garcia-Finana et al., 2003) using the software STEPanizer (Tschanz et al., 2011). The entire span of the OB (7 to 9 coronal sections separated by 40 μm) was evaluated by overlaying a 400-point grid for point counting. We used Gundersen’s coefficient of error (CE) to estimate the precision of the stereology method (Gundersen and Jensen, 1987; Gundersen et al. 1999).

### Densitometry

Adobe Photoshop (Adobe, Mountain View, CA) was used to analyze levels of HuC/D and GFAP staining by determining the optical density (OD) of stained samples. For each animal, 2 to 4 sections throughout the OB were quantified and averaged. Images from coronal sections taken at 20x magnification were converted to 8-bit gray scale, and mean luminosity values were obtained in both contra- and ipsilateral sides. Gray values were converted to optical density (OD) by the following formula: OD =-log (intensity of background/intensity of area of interest). Then, the percent change between contra- and ipsilateral sides was calculated.

### Microglia quantification and characterization

Microglial cells (4C4 immunoreactive) were manually quantified. For each animal, 4 to 6 sections throughout the OB were quantified and averaged. Profiles were categorized by morphological features characteristic of zebrafish microglial activation states described as ramified (1A, 2A, 1R, and 2R); transitioning (3A, 4A, 3R, and 4R); and activated (5A, 6A, 5R, and 6R; Jonas et al. 2012; Var and Byrd-Jacobs, 2019).

### Olfactory-mediated behavioral assays

The behavior apparatus consisted of a white cylindrical chamber with 30 cm diameter outfitted with two surgical tubes found on opposite sides about 8 cm above the base. This chamber was filled with 2.5 liters of fresh fish water before each experiment. Attached to the tubes, syringes were used to administer 3mL of odorant solution or vehicle (PBS). An overhead digital camera was set 1 meter above the chamber to capture swimming behavior (Paskin and Byrd-Jacobs, 2015). To minimize the effect of the experimenters’ presence, the chambers were surrounded by white panels with perforations for the odorant tubes. Odorants were administered behind the panels. Fish were fasted for 48 hours in individual tanks before the experiment and were placed individually in the behavior chamber to acclimate for 1 hour (the last 30 minutes in the absence of movement or sound). For each trial, fish were recorded 30 seconds prior to odorant delivery and then exposed simultaneously to odorant solution and PBS on opposite sides of the chamber to discard swimming differences due to changes in water flow. The recording continued for 30 seconds after odorant delivery. After each trial, fish were transferred to another chamber with fresh water and left to acclimate for at least 60 minutes until the next trial. Fish underwent 3 or 4 behavioral trials per day. For each trial, fish were exposed to 3 ml of 100 μM solutions of alanine, urea with taurocholic acid (TCA), or cadaverine solutions (all from Sigma-Aldrich) in PBS. The odorant tube was kept the same to control for possible leftover solution in the next trial, but the side of the odorant delivery was randomized for each trial.

### Video Analysis

All the swimming behaviors we assessed correspond to well-characterized, stereotypical behaviors described extensively in the literature (Kalueff et al., 2013). An animal tracking software, ToxTrac (Rodriguez et al., 2018), was used to analyze swimming behavior recordings after alanine and urea + TCA exposure. Analyses were divided into the 30 seconds of pre-odorant delivery and the 30 seconds of post-odorant delivery. Each recording was uploaded to iMovie in order to increase the contrast to allow for better detection on ToxTrac. Swimming speed (mm/s), distance swam (mm), and number of directional changes were obtained from ToxTrac. For cadaverine analyses, we coded time (in seconds) spent in normal swimming, erratic swimming, and freezing. Erratic swimming was characterized as a sudden increased speed of movement and rapid, sharp directional changes. Absence of swimming in between normal and/or erratic swimming was characterized as freezing. We calculated the percent change of response in pre- and post-trials using the formula: percent change= (pre-trial value – post-trial value) / post-trial value × 100 or the percent time spent in erratic swimming or freezing.

### Statistical analyses

Comparisons between groups were carried out using Analysis of Variance (ANOVA) with Tukey’s post hoc tests. P-values less than 0.05 were considered significant. Behavioral data were analyzed using a One-Sample Wilcoxon test with a baseline of 100% to compare the percentage change of response among groups. All statistical analyses were performed using GraphPad Prism Software (GraphPad, San Diego, CA).

