## Supplemental figures for "Structural regeneration and functional recovery of the olfactory system of zebrafish following brain injury"

| | Average volume ($mm^3$) | | Gundersen's coefficient | |
|---|---|---|---|---|
| | contra | ipsi | contra | ipsi |
| ctrl | 0.0293 | 0.0284 | 0.025 | 0.024 |
| 1 dpl | 0.0299 | 0.0198** | 0.019 | 0.026 |
| 21 dpl | 0.0311 | 0.0285 | 0.013 | 0.014 |

***Supplemental table 1. Effects of an excitotoxic lesion on olfactory bulb 's volume.***
*Average OB volume and Gundersen's coefficient of volumes obtained from semi-serial H&E-stained coronal sections (n = 3-6). One-way ANOVA. **p < 0.001.*

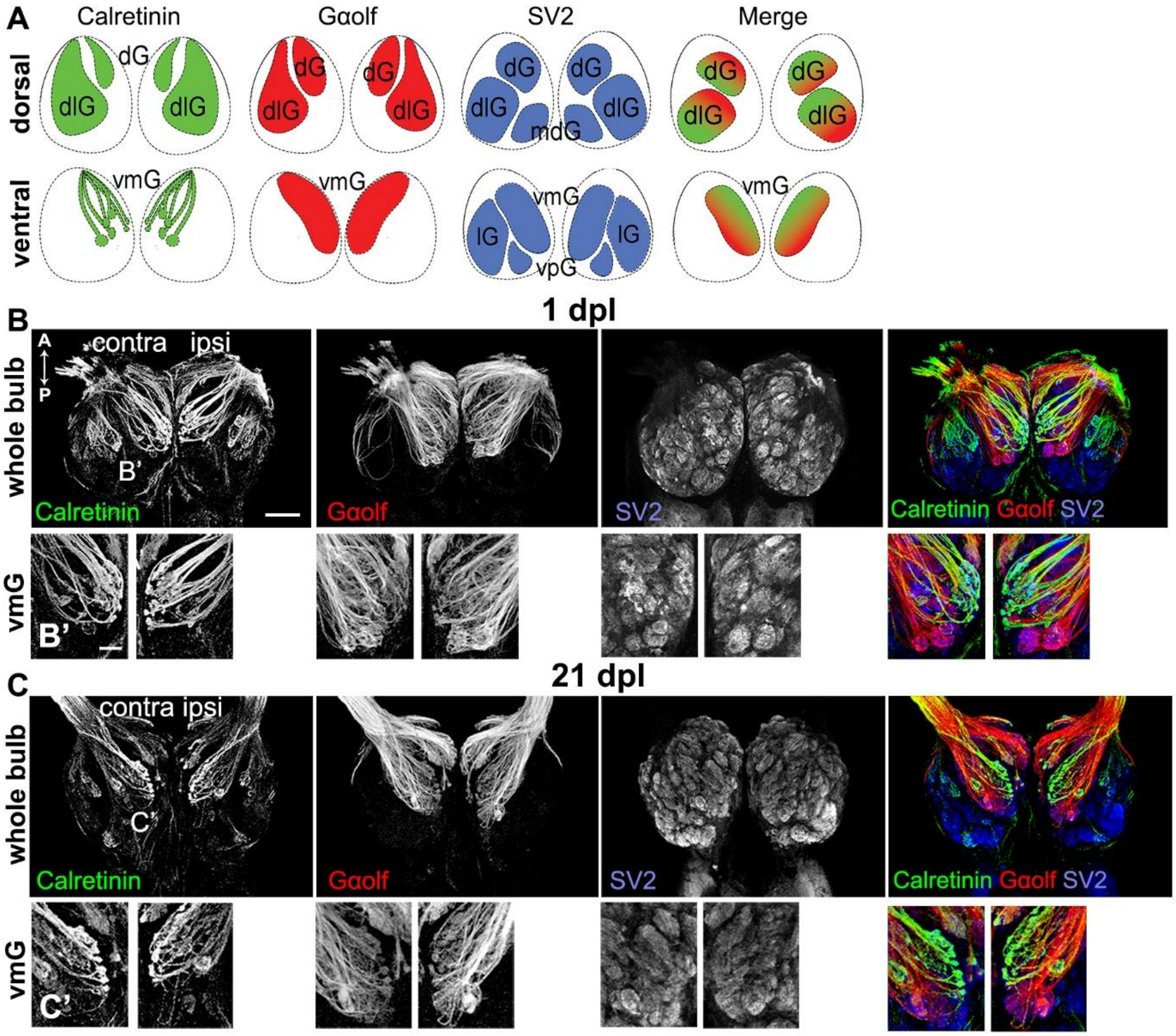

***Supplemental figure 2. Effects of QA lesion on olfactory glomerular morphology.***
*A) Schematic of the dorsal and ventral OB clusters analyzed. The ventral views are inverted to show the ipsilateral side on the right. Abbreviations are as follows: dG, dorsal; dlG, dorsolateral; mdG, mediodorsal lG, lateral; vmG, ventromedial; vpG, ventroposterior.*
*B) and C) Inverted ventral views (ipsilateral side on the right) of whole-mount OB of 1-dpl (B) and 21-dpl (C) groups immunostained against calretinin (left panels), Gαolf (middle panels), and SV2 (right panels). Scale bar: 100 µm.*
*B') and C') Magnified views of the ventromedial cluster from (B) and (C). Green: calretinin; red: Gαolf; blue, SV2. Scale bar: 20 µm.*

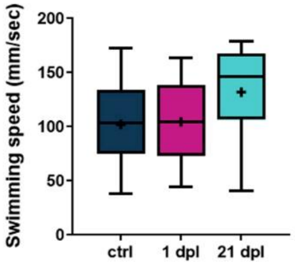


***Supplemental figure 3. Effects of QA bulbar lesion on swimming behaviors.***
*Quantification of swimming speed in ctrl, 1-dpl and 21-dpl fish (n = 8-10). Box plots indicate mean (+), quartiles (boxes) and range (whiskers). One-way ANOVA, not significant.*

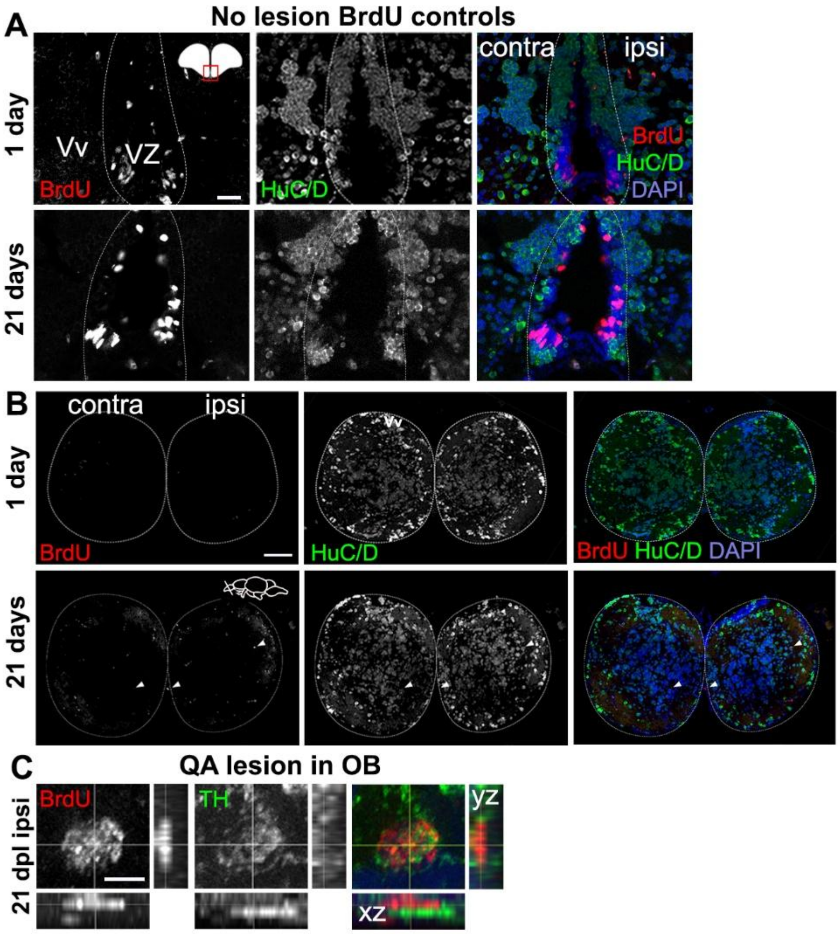


***Supplemental figure 4. Cellular proliferation in the telencephalon and the OB.***
*A) Double immunohistochemistry of BrdU (left panels) and HuC/D (middle panels) in the neurogenic niche of the VZ (ventricular zone, indicated by dashed lines) and the Vv of unlesioned fish 1 day (top panels) and 21 days (lower panels) post-BrdU exposure. Red: BrdU; Green: HuC/D; Blue: DAPI. Scale bars: 20 µm.*
*B) Double immunohistochemistry of BrdU (left panels) and HuC/D (middle panels) in the OB of unlesioned fish 1 day (top panels) and 21 days (lower panels) post-BrdU exposure. Red: BrdU; Green: HuC/D; Blue: DAPI. Scale bars: 20 µm.*
*C) Orthogonal projections (xz, yz) of magnified views from the ipsilateral side of a 21-dpl OB showing a newborn periglomerular BrdU+/TH+ cell. Scale bar: 10 µm.*
